# Cortical Tension Initiates the Positive Feedback Loop Between E-cadherin and F-actin

**DOI:** 10.1101/2021.02.23.432578

**Authors:** Qilin Yu, William R. Holmes, Jean P. Thiery, Rodney B. Luwor, Vijay Rajagopal

**Affiliations:** Department of Mechanical Engineering, University of Melbourne, Australia; Department of Physics and Astronomy, Vanderbilt University, Nashville, Tennessee, United States of America; Department of Surgery, The University of Melbourne, The Royal Melbourne Hospital, Parkville, Australia; Department of Biochemistry, Yong Loo Lin School of Medicine, National University of Singapore, 8 Medical Drive, MD7, #02-03, Singapore, 117597, Singapore; Guangzhou Institute of Biomedicine and Health, Chinese Academy of Science, Guangzhou, People’s Republic of China; CNRS Emeritus CNRS UMR 7057 Matter and Complex Systems, University Paris Denis Diderot, Paris, France; INSERM UMR 1186, Integrative Tumor Immunology and Genetic Oncology, Gustave Roussy, EPHE, PSL, Fac. de Médecine - Univ. Paris-Sud, Université Paris-Saclay, 94805, Villejuif, France

**Author notes:** **Corresponding Author:** Vijay Rajagopal, **Email:**. **Author Contributions:** Q.Y., V.R., R.B.L. designed research; Q.Y. performed research; Q.Y. V.R. analyzed data; and Q.Y., W.R.H., J.P.T., V.R. wrote the paper. **Competing Interest Statement:** The authors declare no competing interest. **Classification:** Major: BIOLOGICAL SCIENCES; Minor: Biophysics and Computational Biology.

**Keywords:** cadherin actin interaction, cell adhesion, intracellular signaling

## Abstract

Adherens junctions (AJs) physically link two cells at their contact interface via extracellular homophilic interactions between cadherin molecules and intracellular connections between cadherins and the actomyosin cortex. Both cadherin and actomyosin cytoskeletal dynamics are reciprocally regulated by mechanical and chemical signals, which subsequently determine the strength of cell-cell adhesions and the emergent organization and stiffness of the tissues they form. However, an understanding of the integrated system is lacking. We present a new mechanistic computational model of intercellular junction maturation in a cell doublet to investigate the mechano-chemical crosstalk that regulates AJ formation and homeostasis. The model couples a 2D lattice-based model of cadherin dynamics with a continuum, reaction-diffusion model of the reorganizing actomyosin network through its regulation by Rho signaling at the intercellular junction. We demonstrate that local immobilization of cadherin induces cluster formation in a cis less dependent manner. We further investigate how cadherin and actin regulate and cooperate. By considering the force balance during AJ maturation and the force-sensitive property of the cadherin/F-actin linking molecules, we show that cortical tension applied on the contact rim can explain the ring distribution of cadherin and F-actin on the cell-cell contact of the cell-doublet. Meanwhile, the positive feedback loop between cadherin and F-actin is necessary for maintenance of the ring. Different patterns of cadherin distribution can be observed as an emergent property of disturbances of this feedback loop. We discuss these findings in light of available experimental observations on underlying mechanisms related to cadherin/F-actin binding and the mechanical environment.

**Significance Statement:** The formation, maintenance and disassembly of adherens junctions (AJs) is fundamental to organ development, tissue integrity as well as tissue function. E-cadherins and F-actin are two major players of the adherens junctions (AJs). Although it is well known that cadherins and F-actin affect each other, how these two players work together to maintain the intercellular contact is unclear. Using a novel mechano-chemical model of E-cadherin and F-actin remodeling, we demonstrate that a positive feedback loop between cadherins and F-actin allows them to stabilize each other locally. Mechanical and chemical stimuli applied to the cell adhesion change E-cadherin and F-actin distribution by consolidating or interrupting the feedback loop locally. Our study mechanistically links mechanical force to E-cadherin patterning at cell-cell junctions.

## Main Text

### Introduction

Adherens junctions (AJ) are protein complexes in localized plasma membrane domains that anchor neighboring cells by linking to the actin cytoskeleton [1]. As a major component of the AJ, E-cadherin is a transmembrane protein that mediates the links between the two cells and works as a core regulator involved in morphogenesis, tumorigenesis and cancer progression [2, 3]. Expression and dynamics of E-cadherin have a direct impact on the mechanical integrity of tissues as well as intracellular signaling pathways, such as TGF-β, Wnt and EGF [4, 5]. Several key components to the formation and homeostasis of adherens junctions have been identified in recent years.

Classical cadherin mediates adhesion through the binding of the extracellular region of cadherins from opposing cells in a calcium-dependent manner, called cadherin trans-dimer. Cadherin proteins from the same cell interact with each other through cis interaction. As two cells are brought into contact, cadherin monomers from opposite membranes start to interact and form trans-dimers. The proximity of two cells differentiates the contact region from the rest area, or contact-free area, since the membrane curvature prevents cadherins on the rest area of the opposing cells to reach each other. Wu. et al proposed that cell-cell contact acts as a trap for E-cadherin trans dimers by limiting their diffusion, increasing E-cadherin concentration in the contact area. They termed this mechanism of E-cadherin clustering as a diffusion-trap. The cooperation between cis and trans in cadherin cluster formation brings more cadherins from the rest area to the contact region, further consolidating cell-cell contact [6].

Upon the proximity of two cells, the interactions between cadherin on the opposite membranes activate a series of signaling cascades [7], which triggers reorganization of the actin cytoskeleton and breaks the force equilibrium at the intercellular contact [8–10]. The intercellular contact evolves until a new force equilibrium is achieved. Members of Rho GTPase family, including Rho, Rac, are believed to play pivotal roles during the maturation process of the intercellular junction. Rac, an initiator of the actin branching network and lamellipodia, is upregulated rapidly once the cells touch each other, and is blocked if the cadherin binding sites are inhibited [11–15]. On the other hand, Rho, a regulator of myosin, which contracts and stiffens the actin network, is down-regulated by cadherin ligation through the cross-talk between Rho and Rac [16].

Notably, cadherin dynamics is also regulated by the cell cortex right beneath the membrane, which is mainly composed of actin and actin-binding proteins. The actin network impedes cadherin diffusion primarily in two ways: (i) delimiting E-cadherin with F-actin cortical meshwork [17]; (ii) binding to E-cadherin via linking molecules such as α-catenin, vinculin and EPLIN [18–20]. Experimental studies indicate that E-cadherin, β-catenin and α-catenin form a minimal cadherin-catenin-complex (CCC) that binds to actin cytoskeleton [19]. In comparison with the binding affinity between αE-catenin and E-cadherin/β-catenin (*K_d_* ~ 1nM) [21, 22], αE-catenin binds to F-actin with a much lower binding affinity (*K_d_* < 1μM) [23–25]. Thus, the binding between αE-catenin and F-actin is a deterministic factor for the interaction between E-cadherin and actin cytoskeleton. Interestingly, this binding is also force-sensitive and acts as a two-state catch bond, which reaches the maximum lifetime when the force sustained in the bond is around 8 pN [19].

X-dimer, one of two conformations of the cadherin trans-dimer found in the extracellular domain of the contact interface, can also explain the force-sensitive property of AJ. The force subjected by each trans-dimer under resting conditions is around 5 pN [26]. However, X-dimer acts as a catch-slip bond and reaches the longest lifetime when the force sustained in each X-dimer is around 25-35 pN [27], ensuring the capability of AJ to resist external forces.

Recently, several computational approaches have focused on cadherin dynamics [9, 25, 26]. These studies shed light on cooperation between trans and cis, cadherin structure, and how the membrane environment affects cadherin dynamics. However, roles of the mechanical environment and cytoskeleton remodeling on cadherin dynamics and intercellular contact formation are yet to be investigated. We present a new computational model of adherens junction formation and maturation at the contact interface of a cell doublet. We use the model to study the crosstalk between mechanical forces, and biochemical signals that regulate E-cadherin and actin dynamics and discuss the implications of the model in light of experimental findings [17, 28, 29] (see [30, 31] for a review of similar approaches).

A previous computational study suggested that upon cell-cell contact, E-cadherin tends to move from the free contact area to the contact region due to the trans-dimer diffusion trap [6]. Here, by adapting and implementing the same lattice-based model, we demonstrate immobilization of E-cadherin in the contact region through actin binding locally inducing cluster formation, even with very weak cis binding affinity (0 - 2 k_B_T). Moreover, we observe cluster formation exclusively in the immobilization trap. when we simulate trans-dimer diffusion trap and immobilization trap simultaneously.

During intercellular contact formation between two cells in suspension, cadherin and actin diminish at the contact center, and accumulate to the contact edge gradually and form a ring-like distribution. This has been observed in several cell types, including Madin-Darby canine kidney cells (MDCKs), carcinoma cells, and zebrafish epiblast [28, 32–34]. To unravel the physical principle of the ring distribution formation of E-cadherin and actin on cell-cell contact of a cell-doublet, we coupled the lattice-based model of E-cadherin dynamics to a minimal reaction-diffusion model of the actomyosin network. We tested the hypothesis that the difference in E-cadherin mobility between the contact center and the contact rim triggers the ring distribution formation. The difference in mobility comes from the cortical tension applied on the contact rim and the force-sensitive feature of E-cadherin/F-actin linking proteins, such as α-catenin and vinculin. Model simulations also imply the necessity of the positive feedback loop between E-cadherin and F-actin to maintain the ring distribution. Furthermore, we investigate how different disturbances to the feedback loop modulate E-cadherin and F-actin distribution and reproduce experimental observations [3, 8, 27].

## Computational Model

### Lattice-based model of E-cadherin clustering dynamics

To model E-cadherin’s clustering dynamics, we modified and employed a lattice model built previously by Wu *et al.* [6]. As shown in Fig. 1 B the side length of each lattice site is 7 nm, which is the interval between E-cadherin molecules in the crystal structure [35]. A and B represent monomers in the two opposing cell membranes, and D represents the trans-dimer, A + B ⇄ D.

**Figure 1.**
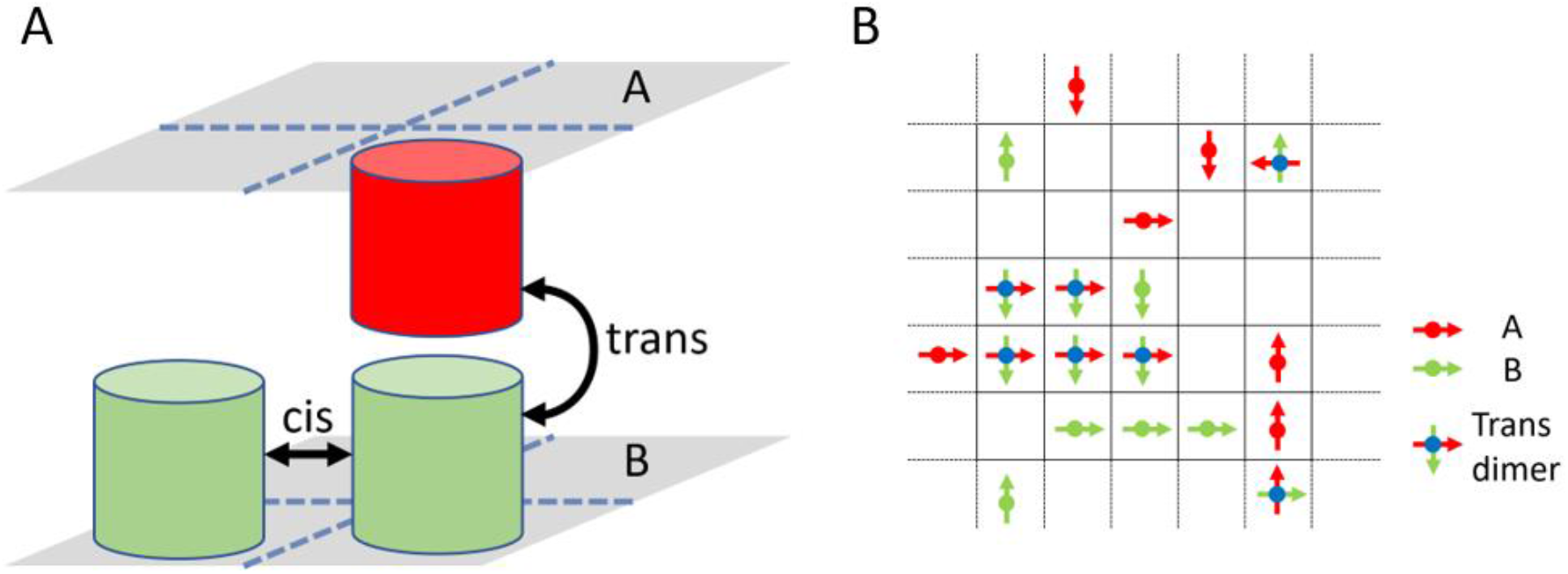
(A) Schematic illustration of E-cadherin interactions. (B) The 2D field composed of square lattices, representing the contact area of a cell doublet. Each lattice can only be occupied by one monomer from each cell. Occupancy by a monomer A from the upper membrane (red arrow) and a monomer B from the bottom membrane (green arrow) simultaneously is necessary for the formation of a trans-dimer. Monomers from the same surface can interact with each other through cis-interaction.

In the model, double occupancy of one lattice site by two As or two Bs is not allowed. However, A and B must be placed in one lattice site to form a trans-dimer. Each particle may orient toward any of the four possible directions, N, E, S and W. Thus, there are four types of monomers on each side, A_N_, A_E_, A_S_, A_W_, and B_N_, B_E_, B_S_, B_W_ (Fig 1 B). Consistent with the crystal structure, a trans-dimer can only be formed by an A and a B, where B’s orientation is rotated clockwise by 90° relative to A’s orientation (e.g. D_NE_ = A_N_ - B_E_). Two monomers on the same surface can interact via their cis-interfaces only if they occupy the nearest-neighbor sites along the same axis, e.g. A_N_ – A_N_. A trans-dimer can only interact with the same type of trans-dimer, e.g. D_NE_ – D_NE_. However, due to the two orthogonally oriented monomers in a dimer, a trans-dimer can interact with dimers and monomers in both axes.

The length of a single time step used in the simulation can be calculated in two dimensions, where Δ*t* is the time step; *x* is the distance between two lattices; D is the diffusion coefficient of an E-cadherin in 2D.

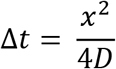

We defined all monomers and dimers that are not in cis-interactions as free particles. In each time step of the simulation, all free particles are randomly selected to attempt a step movement. Displacement of particles to occupied sites will be rejected. After the movement, A and B in the same lattice can form a transdimer with probability γ/ (1+ γ) where γ = exp[Δg^0^_trans_/k_B_T]. Δg^0^_trans_ represents free energy change of trans interaction in the lattice-based model, which is derived from the molar binding free energies measured in 3D solution. Additional details of the calculation can be found in Wu’s work [6]. Two monomers in a dimer will dissociate with the probability 1/ (1+ γ). Cis interactions behave similarly in the model, except that the reactants are monomers from the same surface, meaning two As or two Bs.

### Coupling the lattice-based model with the Reaction-Diffusion model to simulate intercellular contact formation

E-cadherin and F-actin rings have been observed in several cell types, including Madin-Darby canine kidney cells (MDCKs), carcinoma cells, and zebrafish epiblast [28, 32–34]. We hypothesize that the ring distribution results from the feedback loop between actin remodeling and cadherin dynamics. In our model, cadherin ligation upregulates local actin polymerization in a Rac-dependent manner and cortical actin microfilaments inhibit cadherin diffusion via binding (Fig. 2 C). The mechanical environment on the contact can regulate E-cadherin/F-actin binding via mechanosensitive linking proteins, which then controls cadherin diffusion. Here, we introduce an integrated model that couples the lattice-based E-cadherin model with a simplified reaction-diffusion (RD) finite element model of F-actin and related regulatory proteins (Fig. 2 C) to test our hypothesis.

**Figure 2.**
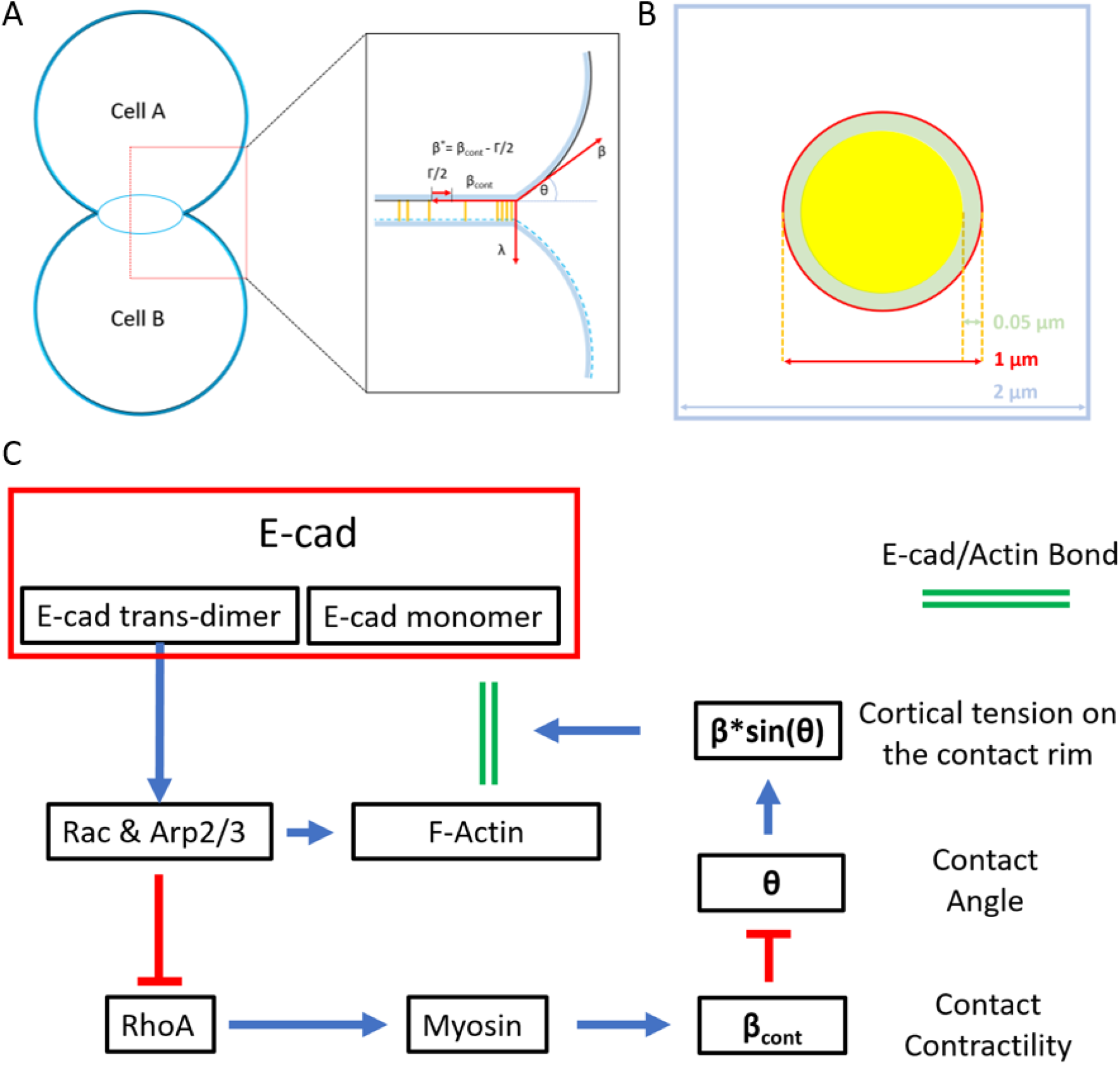
Illustration of the integrated model (A) 3D representation of a cell doublet with one cell stacked on the other cell. Inset: Cell membrane is in black and actin cortex is in blue. Tension analysis was shown only on the upper cell to improve visualization. At the equilibrium state, tensions on the contact rim will be balanced by each other. Tensions at the contact rim includes cortical tension, β, at the free cell surface; tension on the contact β^*^, which equals to the tension from the remaining actin cortex, β_cont_, minus adhesion tension Γ/2; the linking tension λ, sustained in the cadherin trans-dimers (orange line between two cells) and the force-sensitive molecules such as vinculin and α-catenin. The contact angle θ, and the linking tension λ, is determined by the relationship between β and β^*^. (B) Schematic illustration of the 2D simulation area of the coupled model. The region inside blue square and outside red circle represents the free contact area, where trans-dimer cannot be formed. Area in the red circle represents the cell-cell contact region, where the trans-dimer formation is allowed. Within the contact region, green circular ring with a width of 0.05 μm represents the contact edge, and yellow circle represents the contact center with a radius of 0.45 μm. (C) Signaling pathways applied in the model. Formation of trans-dimers upregulates Rac, which works as an upstream regulator for F-actin polymerization. Rac also inhibits RhoA, myosin and contractility of cell cortex beneath the contact membrane via p190B-RhoGAP. The change of force balance on the intercellular contact leads to an increment of the contact angle θ and a larger tension sustained in the bonds between E-cadherin and F-actin. A force around 10 pN maximizes the lifetime of the E-cad/F-actin linkage due to the force sensitive bonds between α-catenin/vinculin and F-actin.

We test our model hypothesis against previous experimental observations of e-cadherin and actin dynamics in S180 cell doublets [28, 29]. In these experiments, two S180 cells were suspended within a microfluidic device and e-cadherin and actin distributions at the contact rim were observed. We compared our integrated model of the contact interface primarily against these data. In the new model, the lattice field is composed of 286 × 286 lattices, a 2μm by 2μm region with periodic boundaries (Fig. 2 B). The central circular area with 1μm diameter represents the intercellular contact where the trans-dimers can be formed. The region’s diameter used in the model is scaled down approximately ten times with respect to the intercellular contact area of the S180 cell doublet to reduce computational cost. All the lattice-sites outside the central area represent the free contact area, where trans-dimers cannot be formed (Fig. 2 B).

A corresponding simplex triangulated mesh is constructed to solve the PDEs of signaling pathways incorporated in the new model using the finite element method (Fig S5). Details of the process of mesh building and the incorporated signaling pathways can be found in supplemental materials.

Within each time step of the RD model, the state of each E-cadherin monomer in the lattice-based model is updated to the finite element mesh of the RD model. Upon stimulation by E-cadherin homophilic interactions, the dynamics of species involved in Rho, Rac and Cdc42 are solved based on the RD equation:

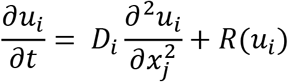

where u_i_ stands for the local concentration of chemical species i, which is a function of time, t, and distance x_j_ in dimension j; D_i_ is the diffusion coefficient of the species; R(u_i_) accounts for all local reaction kinetics of chemical species i.

E-cadherin is anchored once it is trapped by the actin network and becomes mobile only after the bond dissociates. The bond between E-cadherin and F-actin is simplified as a direct interaction in the RD model, since the bond between α-catenin and β-catenin/E-cadherin is much stronger than the bond between α-catenin and F-actin. The association rate between E-cadherin and F-actin is held constant. However, the dissociation rate is a function of the link tension due to the force-sensitive nature of the bond between α-catenin and F-actin (Fig. S7).

The forces involved at the intercellular contact are illustrated in Fig. 2 A and include: (i) cortical tension from the actin cortex beneath the free contact membrane β;(ii) tension from the remaining actin cortex beneath the contact membrane β_cont_; (iii) total binding energy of cadherins on the cell-cell contact, or the adhesion tension Γ (Γ/2 per cell); (iv) and the link tension λ, sustained in the trans-dimers [10]. β_cont_ is determined by the total amount of activated myosin (see supplementary materials), and Γ is calculated based on the total number of trans-dimers on the contact region. Given a constant cortical tension β, we can calculate the contact angle θ, and the link tension λ by solving the force equilibrium equation:

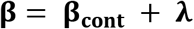

Cortical tension β is held constant at 4000 pN/μm [10]. Effects of cortical tension β on the E-cadherin dynamics are studied in Figure S3. The initial value of β_cont_ is set same as β. The contact angle θ is 0° at the beginning when two cells come into close proximity and increases as actin and myosin networks in the central contact degrade during the contact maturation process. In all studies using the coupled model, it is assumed that the cortical tension β is only applied in the 0.05 μm thick edge of the contact region, roughly 1/10 of the thickness of the actin cortex in the cell [36, 37], green circular ring in Fig. 2B. All chemical species are homogeneously distributed on the contact at the start of the simulation. As E-cadherin monomers from opposite membranes meet each other and form trans-dimers, the feedback loop between E-cadherin and F-actin is triggered.

## Results

### Cooperation Between Contact-induced “trans-dimer diffusion trap” and F-Actin-related “immobilization trap”

As soon as two suspended cells touch each other, the contact area increases and reaches a steady state in minutes. During the maturation process, E-cadherin diffuses from the free contact membrane to the contact area because the trans-dimer formation is limited to the contact area [6]. We refer to this effect as the contact-induced “trans-dimer diffusion trap”. “trans-dimer diffusion trap” applies only to the membrane regions where the distance between two opposite membranes is smaller than the maximum distance for a trans-interaction to occur.

On the other hand, at an intercellular contact, E-cadherin can be trapped by the actin cytoskeleton through linking proteins, such as α-catenin and vinculin. Experimental results show the alignment between E-cadherin clusters and F-actin [38]. We propose that due to the force-sensitive property of α-catenin and vinculin, E-cadherin’s diffusivity could be different in different cell-cell contact regions in response to the local cytoskeletal and force distribution. Locally lower diffusivity serves as another type of trap. We define this mechanism as F-actin related “immobilization trap”.

To evaluate the effects of the two mechanisms on clustering, we simulated E-cadherin dynamics on a lattice field composed of 50 × 50 lattice sites with periodic boundaries (Fig. 3 A-D). All simulations in Fig. 3 are carried out with a concentration of 800 monomers/μm^2^ on each surface, a biological meaningful concentration according to literature or 0.04% of the occupied lattices [6, 17, 39]. To demonstrate the effect of two mechanisms, we designed four sets of studies with a broad range of Δg^0^_cis_ and Δg^0^_trans_ and analyzed the maximum cluster size in each simulation (Fig. 3 E-H).

**Figure 3.**
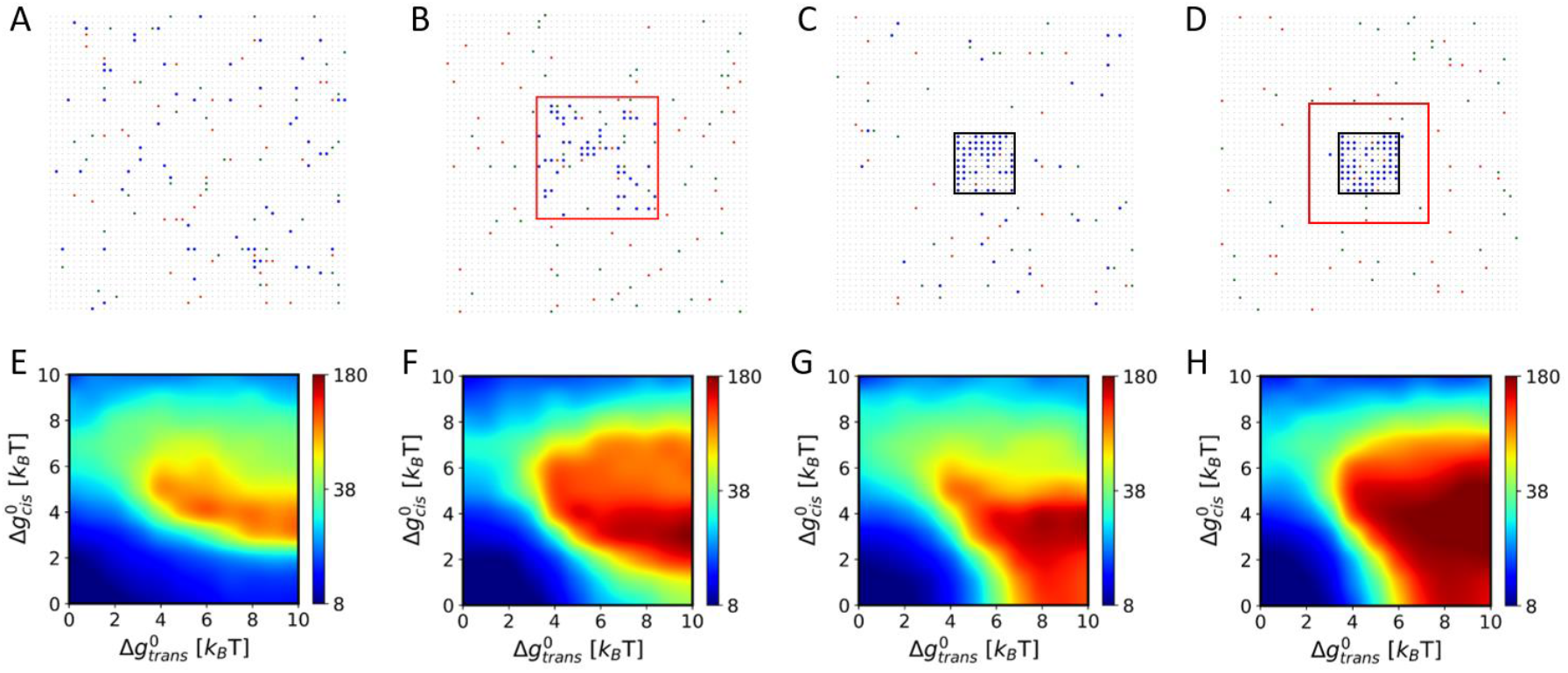
E-cadherin dynamics was simulated on a region with 50 × 50 lattice sites, with the periodic boundaries. In all cases, the monomer concentration is 0.04 on each side. Red and green dots represent E-cadherin monomers on each side of the cell-cell contact respectively. Blue dot represents trans-dimer. Snapshots in a-d were taken from the simulations when the maximum cluster appears in the simulation with 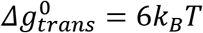, 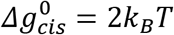 Heatmaps in E-H show the maximum cluster size in each simulation with different 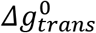. and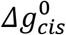 values, from 0kT to 10kT. Each data point represents the mean value of the maximum cluster size of 5 simulations. In A, E, trans-dimers can be formed everywhere. All E-cadherin monomers and trans dimers can diffuse freely on all the lattices. In B, F, trans-dimers can only be formed in the central 20 × 20 lattice sites (Red square), which refers to ‘trans-dimer formation trap’ In C, G, trans-dimers can be formed everywhere, but will be immobilized once they are in the central 10 × 10 lattices (Black square), and can only leave after the trans dimer break down. We define this mechanism as the ‘diffusion trap’ In D, H, “trans-dimer formation trap” is applied to the central 20 × 20 lattices (Red square). “immobilization trap” is applied to the central 10 × 10 lattices (Black square).

In the first set of studies (Fig. 3 A, E), trans-dimers can be formed everywhere. Monomers and trans-dimers can freely diffuse on all the lattice sites. In Fig. 3 E, a sharp boundary can be seen at around Δg^0^_cis_ = 2-6 k_B_T, indicating that large clusters can only be detected in simulations with an intermediate cis-binding affinity. Our findings agree with the results obtained from a previous Monte Carlo study [6, 40]. If Δg^0^cis is under 2 k_B_T, cis interaction is too weak to aggregate E-cadherin proteins together. On the other hand, if Δg^0^_cis_ is above 6 k_B_T, there are many more cis-interactions that limit the number of freely diffusing monomers that would otherwise aggregate into a larger cluster. Apart from the cis-dominating range (Δg^0^_cis_ > 6 k_B_T) and cis-negligible range (Δg^0^_cis_ < 2 k_B_T), 2-6 k_B_T of Δg_0_cis is the range where trans-affinity and the cooperation between trans and cis can occur effectively to form E-cadherin clusters.

In the second set of simulations, we studied the effect of the trans-dimer “diffusion trap”. E-cadherin trans-dimers can only be formed on the central contact region (demarcated with a red border), composed of 20 × 20 lattice sites (Fig. 3 B). The region outside the red border represents the contact-free area where trans-dimers cannot form. Therefore, movements of trans-dimers toward lattice sites outside the constrained region will be rejected. In other words, E-cadherin can only leave the trap as a monomer when the trans-dimer breaks. Based on the comparison between Fig 3 E and F, the boundary expands on the Δg^0^_cis_ axis but not on the Δg^0^_trans_ due to the concentration increase in the trans-dimer diffusion trap. However, if we increase the overall concentration without defining a contact region, the boundary expands on both Δg^0^_cis_ and Δg^0^_trans_ axis (Fig. S1 C).

In the third set of simulations, E-cadherin trans-dimers can form everywhere but will be immobilized once they are formed or enter the central 10 × 10 immobilization zone (demarcated by the black border, Fig. 3 C). This zone represents E-cadherin immobilization by the local cortical actin. Any movements of trans-dimers in the immobilization trap zone will be rejected. The heatmap shows that noticeable clusters can be detected in the simulation with Δg^0^_cis_ smaller than 2 k_B_T (Fig. 3 G), which agrees with a recent experimental observation that cadherin binding to F-actin drives cadherin clustering in a cis-independent manner [38]. Furthermore, this finding suggests that F-actin anchoring also largely determines E-cadherin clustering location, since F-actin anchoring lowers the local Δg^0^_cis_ threshold for cadherin clustering.

In the process of a cell-cell contact formation, both mechanisms mentioned above could be active. If we turn on both traps simultaneously in one simulation, the cluster formation boundary expands again due to further increase of E-cadherin concentration in the central 20 × 20 lattice sites (Fig. 3 H, S1 C). Interestingly, the cluster forms exclusively in the immobilization trap, the central 10 × 10 lattice sites (black square) inside the trans-dimer formation trap (red square) (Fig. 3 D). This finding implies the correlation between the local diffusivity and cluster formation site. In comparison with the first set of simulations, the cooperation between the trans-dimer “diffusion trap” and “immobilization trap” facilitates cluster formation and lower the requirement of Δg^0^_cis_ for cadherin clustering.

### Cortical tension initiates a positive feedback loop between E-cadherin and F-actin

As mentioned above, E-cadherin homophilic interactions upregulate actin polymerization by regulating Rho family GTPases. Meanwhile, F-actin regulates E-cadherin’s dynamics locally and induces cluster formation. This positive feedback loop allows F-actin and E-cadherin to strengthen each other. Both F-actin and E-cadherin are homogeneously distributed at the contact region at the beginning of contact formation but accumulate on the edge as the adhesion matures [Fig. S2 A]. Both α-catenin/F-actin and vinculin/F-actin bonds are sensitive to force with a force of about 8-10 pN, minimizing their lifetime. Thus, we test the hypothesis that cortical tension on the edge initiates a positive feedback loop by increasing the CCC/F-actin bond’s lifetime, assuming that cortical tension is only present at the contact edge.

The distribution of E-cadherin predicted by the model are shown in Fig. 4A for a range of values of parameters representing E-cadherin trans-dimer induced Rac activation (k1) and the E-cadherin to F-actin binding rate (k5). As the cell contact matures, tension sustained in the CCC and the α-catenin/F-actin bonds located at the contact rim increases, which increases the α-catenin/F-actin bond lifetime and allows F-actin to anchor E-cadherin for a longer time (Fig. 2 A, C). This effect leads to a difference in the E-cadherin diffusivity between the contact center and the contact rim. E-cadherin clustering on the contact rim upregulates F-actin formation locally, further anchoring more E-cadherin proteins on the contact rim (Fig. 4 A).

**Figure 4.**
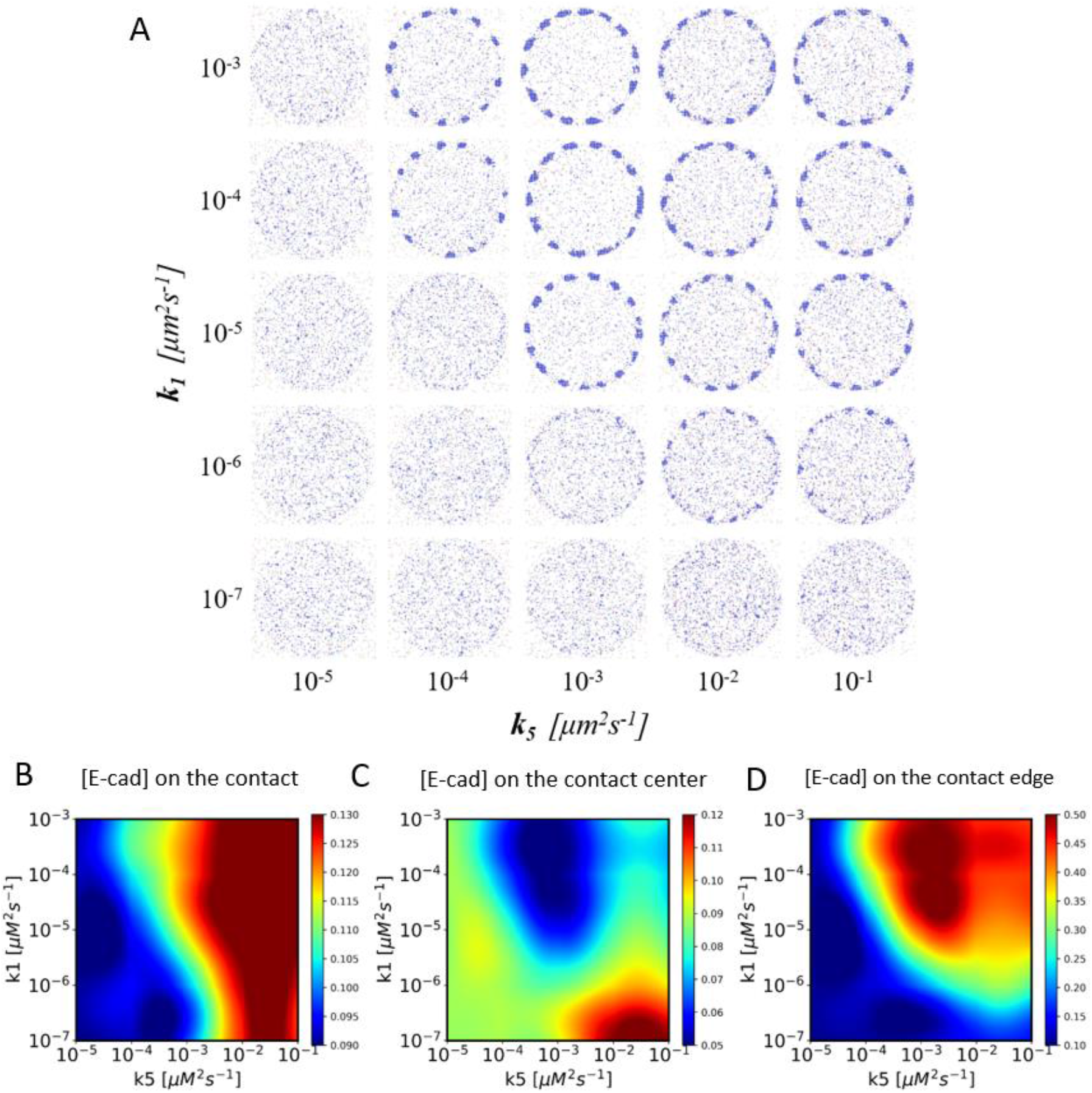
Positive feedback loop between F-actin and E-cadherin matters for contact maturation. (A) Snapshots of the E-cadherin lattice-based model at 300s. Only the circular contact region is shown. E-cadherin concentration is 0.04 on the simulation area all the time. k1 is the trans-dimer related Rac activation rate. k5 is the E-cadherin/F-actin binding rate. 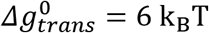, 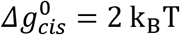. Heat maps of E-cadherin concentration on the (B) whole contact area; (C) the contact center; (D) the contact edge.

See Fig. S2 and Movie S1 & S2 for more details. Simulations with other values of Δg^0^_cis_ and Δg^0^_trans_ show similar E-cadherin and F-actin distribution (Fig. S4). E-cadherin distributions with a wider range of cortical tension and E-cadherin concentration are demonstrated in Fig. S3. Lack of cortical tension on the contact rim leads to failure of the ring’s formation (Fig. S3). These results show that cortical tension initiates this positive feedback loop which brings more E-cadherin and F-actin to the contact rim until cell shape changes stop and a new tension equilibrium is established.

It is known that cells cannot form mature and stable contact without either CCC/F-actin binding [28, 38] or trans-dimerization induced Rac activation and actin recruitment [13–15, 41]. Considering the requirement of both mechanisms for the formation of a mature cell-cell contact, we further evaluated the effects of both reaction rates, trans-dimerization induced Rac activation rate, k1 and E-cad/F-Actin binding rate k5, at wide ranges and analyzed the E-cadherin concentration on the whole contact, contact edge, and contact center respectively (Fig. 4 A, B-D). As expected, E-cadherin rings cannot be formed in simulations with either low k1 or low k5 (Fig. 4 D), indicating that both reactions are necessary to activate the positive feedback loop. As the F-actin-related immobilization trap is discussed in Fig. 3 C, D, G, H, F-actin binding anchors E-cadherin locally, decreasing the local diffusivity and bringing more E-cadherin proteins to the contact region. Effect of this mechanism is consolidated with a larger E-cadherin/F-actin binding rate (Fig. 4 B). Interestingly, with low k1, trans dimerization is not able to upregulate the actin polymerization rate locally to activate the positive feedback-loop and induce ring formation. In conclusion, the model suggests that the difference in the E-cadherin diffusivity caused by the cortical tension on the contact rim is one of the main reasons for the ring formation. Furthermore, the positive feedback-loop between E-cadherin and F-actin is key to maintaining their distribution.

### Variations in F-actin turnover rate may regulate E-cadherin distribution

In cell doublets, one cell can deform and skew to one side spontaneously [28]. Interestingly, E-cadherin, F-actin, myosin networks and other related molecules accumulate to the skewed side of the contact rim, where the contact angle is smaller (Fig. 5 A). The experimental results showed that myosin-regulated actin turnover is the main reason for the asymmetric distribution of E-cadherin and F-actin [28]. We hypothesize that a locally more stable actin network in the skewed side than the opposing side lowers the dissociation rate between CCC and F-actin, decreasing the local E-cadherin diffusivity and bringing more E-cadherin protein to the right side of the contact rim.

**Figure 5.**
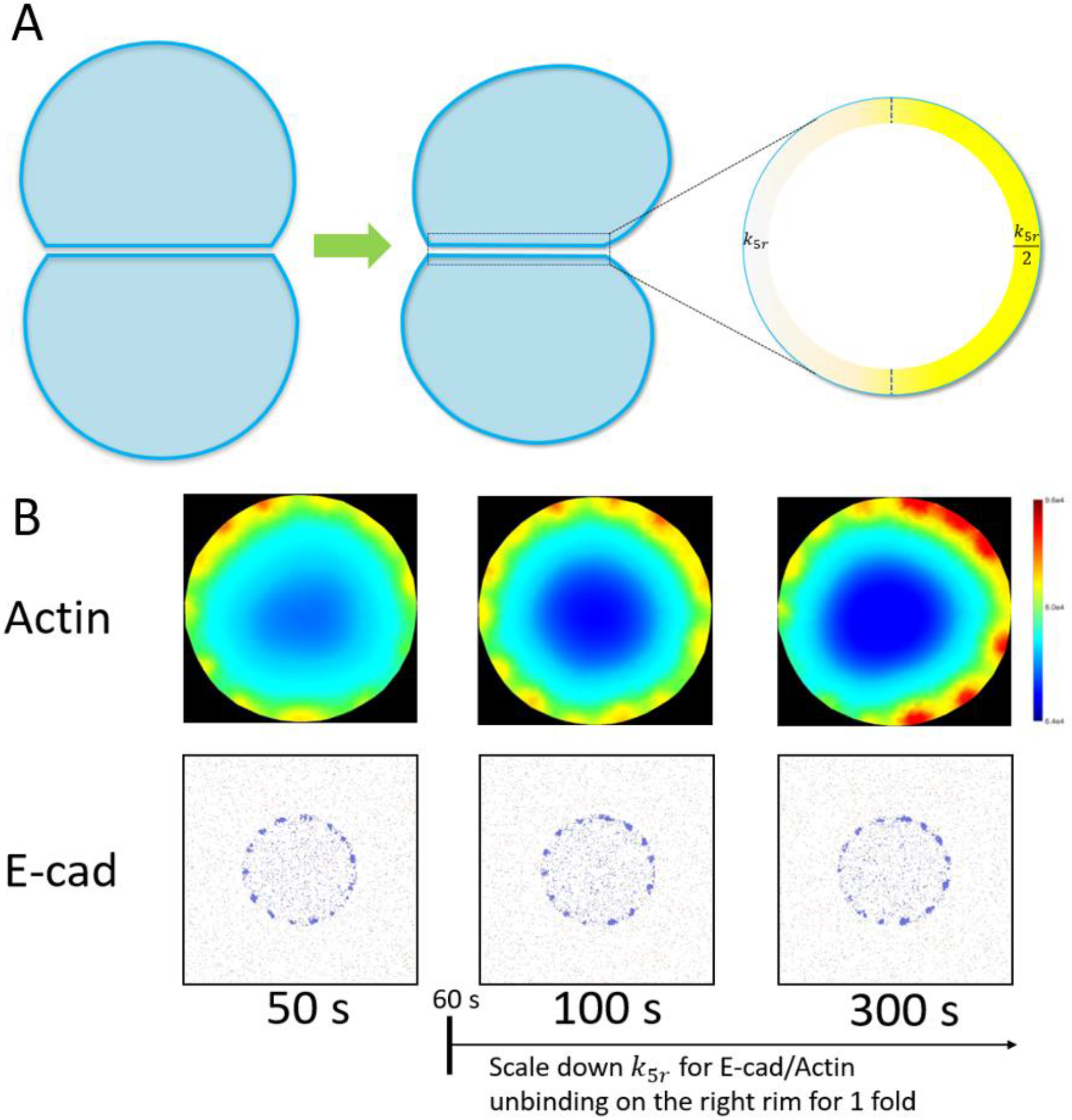
F-actin turnover rate regulates cadherin distribution. (A) Schematics of the deformed cell doublet. E-cadherin molecules accumulate to the skewed side, or the right side. (B) A scaling factor, 0.5, is applied to dissociation rate k5r on the right side of the contact edge after 60s. Both E-cadherin and actin accumulates to the right side of the contact edge due to the locally lower trans dimer diffusivity induced by the more stabilized F-actin network. 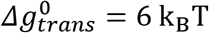, 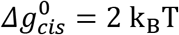 *k*_1_ = 5 ∗ 10^−6^μ*m*^2^*s*^−1^, *k*_5_ = 0.03μ*m*^2^*s*^−1^

To simulate the experimental observation and test the hypothesis of E-cadherin diffusivity, we allow the contact to mature in the first 60s and scale-down the dissociation rate between E-cadherin and F-actin by a “scaling factor” (SF) on the right side of the contact rim, after 60s (Fig. 5 B). E-cadherin gradually moves to the right side of the contact rim and forms clusters there. Although we can still see cluster formation on the left side of the contact rim, the cluster size is much smaller. In this simulation, there are three regions on the cell-cell contact with different E-cadherin diffusivities, D_right-rim_ < D_left-rim_ < D_central-contact_. E-cadherin tends to stay longer on the right side of the contact rim. Due to the positive feedback loop between F-actin and E-cadherin, differences in E-cadherin concentration on these three regions are further consolidated.

### External force regulates E-cadherin distribution by increasing trans-dimer binding affinity

In another study of cell doublets, a flow was introduced cyclically to stretch the bottom cell from the right side (Fig. 6 A) [29]. The deformation of the cell-doublet caused by the flow is similar to the deformation mentioned in Fig. 5, but leads to an opposite effect on E-cadherin and F-actin distribution. E-cadherin proteins accumulate to the stretched side of the contact rim. The force is transmitted to and sustained in the E-cadherin trans-dimers (Fig. 6 A). As introduced above, X-dimer, one conformation of the trans dimer, forms a catch-slip bond and lives longer in the presence of the mechanical stimulation [27]. Thus, we hypothesize that the underlying reason for the asymmetric E-cadherin distribution in this experiment is the locally higher trans-dimer binding affinity due to the stretching.

**Figure 6.**
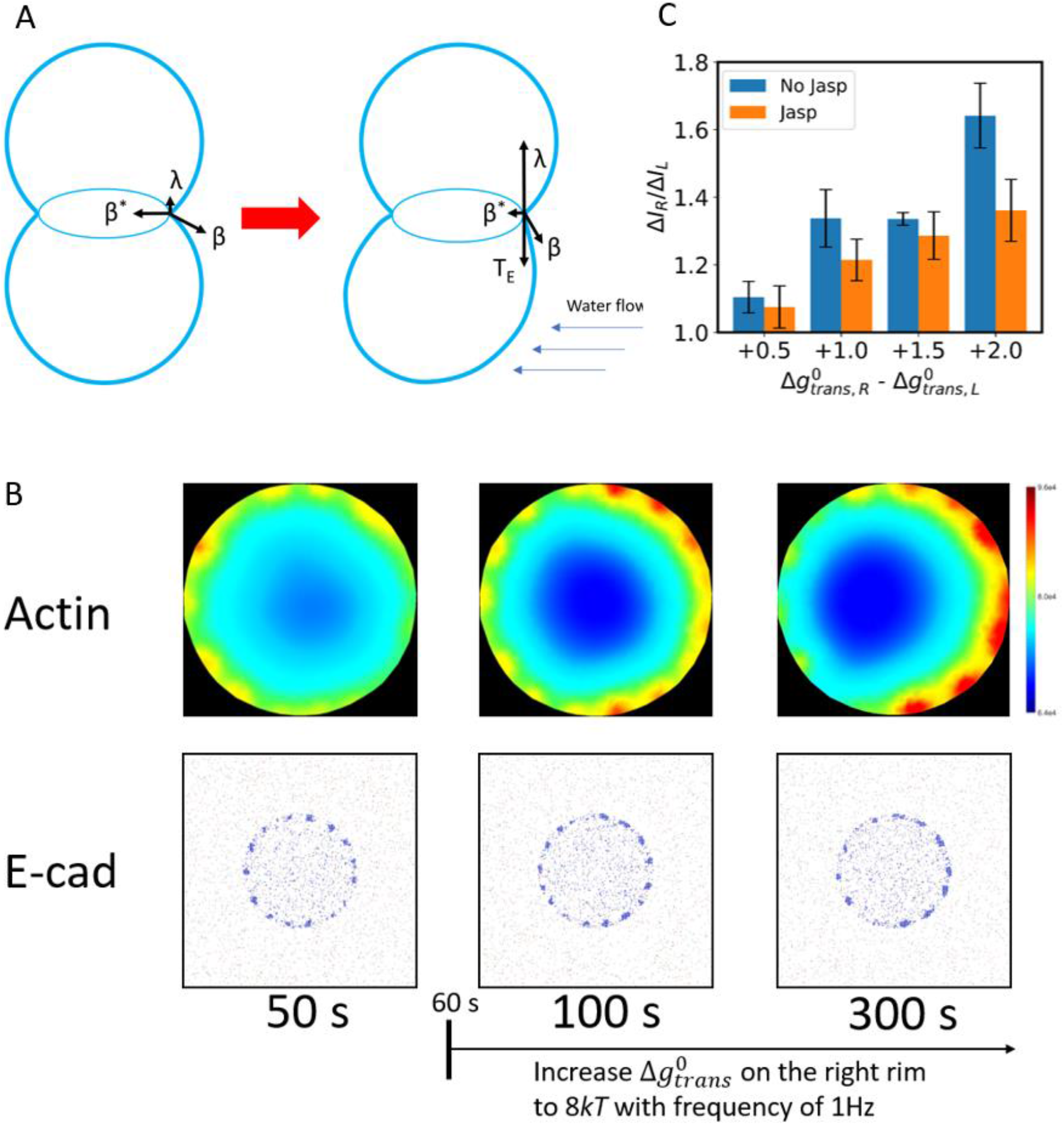
External force induced trans-dimer formation trap and interruption of the positive feedback loop. (A) Tension analysis on the contact rim of the stretched side. Linking tension λ sustained in the E-cadherin trans dimer increases in response to the external stretch. Lifetime of the trans-dimer increases in response to large force. (B) To simulate the effect of the external stretch, 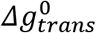 was brought up from 6 k_B_T to 8 k_B_T cyclically with a frequency of 1 Hz after 60s on the right side of the contact rim, and kept 6 k_B_T at all other places. E-cadherin molecules and actin accumulate to the right side of the contact rim gradually. 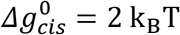, *k*_1_ = 5 ∗ 10^−6^μ*m*^2^*s*^−1^, *k*_5_ = 0.03μ*m*^2^*s*^−1^ (C) The mean ratios between E-cadherin molecules’ concentration change on the right side and left side of the contact rim (ΔI: relative concentration increase). The ratios of simulations with “No jasp” is much higher than the ratios of simulations with “jasp”. The height of each bar comes from the mean value of 5 simulations, and error bars shows the std. The 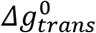 on the right side of the contact rim is increased to 6.5, 7.0, 7.5, 8.0 k_B_T respectively, and kept as 6 k_B_T at all other lattice sites.

To test this hypothesis, we set the Δg^0^_trans_ on the right side of the contact rim to vary between 6 k_B_T and 8 k_B_T cyclically with a frequency of 1 Hz and kept Δg^0^_trans_ at all other places to be 6 k_B_T after 60 s. E-cadherin proteins accumulate to the right side of the contact rim gradually due to the difference in the Δg^0^_trans_, which results in the accumulation of F-actin to the right side (Fig. 6 B). The ring distribution on the contact rim remains, suggesting the co-existence of the two underlying mechanisms, cortical tension inducing ring formation and external force inducing local E-cadherin accumulation.

However, if the actin filament polymerization and stabilization drug, Jasplakinolide (Jasp), is applied to the cell doublet before the introduction of flow, the asymmetric distribution of the E-cadherin disappears when stress is applied to one side of the contact rim in the experimental study [29]. To simulate the effect of the Jasp, we introduced an additional actin polymerization term in the RD model and set the kinetic constant k0 to be a large value, which turns most actin monomers into F-actin. The effect of trans-dimerization on actin polymerization is minimized by doing so. We labeled those simulations with extra actin polymerization as “Jasp”, and the simulation in Fig. 6 C as “No Jasp”.

We analyze model simulation results of the intensity change, *ΔI*, of E-cadherin on the left and right rim before (30-60s) and after (270-300s) a higher Δg^0^_trans_ is applied to the right side of the contact rim. The ratio between *ΔI* on the left and right rim, *ΔI_R_/ΔI_L_*, clearly demonstrated that locally different trans-dimer affinity could induce the asymmetric distribution of E-cadherin and F-actin (Fig. 6 C). However, the interruption of the feedback-loop between trans-dimerization and actin polymerization prevents the accumulation of E-cadherin on the stretched side to strengthen cell adhesion, which might lead to breakdown of cell adhesion. Therefore, our model shows that although external forces induce longer trans-dimer lifetime, which leads to accumulation of E-cadherin locally, disruption of the positive feedback loop can diminish that effect.

## Discussion

Classical cadherin is one of the main players on the intercellular contact, whose concentration and distribution are crucial for tissue integrity. Interestingly, experimental studies showed that cadherin’s distribution is not homogeneous on the contact area and can dynamically vary depending upon cell shape and the external mechanical environment. Here we developed a parsimonious mechanistic model of the interactions between E-cadherin and F-actin in cell doublets to study how feedback between the two influences the dynamic distribution of E-cadherin.

The height of the interfacial shell for the trans-dimer formation is about 10nm. As two cells approaching each other and the extracellular cadherin repeats 1 (EC1) of two cadherins on the opposing membrane are within the interfacial shell, cadherins meet the distance requirement for trans-dimer formation [42]. Using a lattice-based model, we showed that E-cadherin molecules move from the non-contact region to the contact region, which increases the concentration of E-cadherin on the contact region and expands the cluster formation phase boundary. However, membrane curvature driven by membrane fluctuation, cell protrusions facilitated by filopodia and lamellipodia, and size difference of other membrane proteins may prevent E-cadherin molecules in some regions from meeting the distance requirement [40, 43], which could also lead to inhomogeneous E-cadherin distribution at the intercellular contact.

During the maturation process of the intercellular contact, distributions of E-cadherin molecules and F-actin are highly correlated during the process. Moreover, it is known that the actin cytoskeleton regulates cadherin dynamics by anchoring cadherin via linking molecules and delimiting cadherins from reaching each other through actin meshwork. We demonstrated with the new model that spatial variation of E-cadherin mobility can explain the inhomogeneous distribution on the contact zone. Moreover, our results also suggest that the cis-interaction is not indispensable for the cluster formation, which agrees with previous experimental findings [38].

To simulate the maturation process of the cell-cell contact, we incorporated a parsimonious representation of signaling pathways of the Rho family to the lattice-based model of cadherin dynamics. Our results show that the longer lifetime of the cadherin/F-actin linkage induced by the cortical tension can lead to the accumulation of cadherin molecules on the contact edge because of their spatially varying diffusivity on the contact region. Free energies of trans and cis interactions are modulated by other factors, such as calcium level, and mechanical forces. Our results demonstrate that a small change of the binding affinity will not affect the role of cortical tension. However, due to the simplification of the model, effects of intermediary components of signaling pathways that are involved in AJ formation have not been included in the model. F-actin and myosin networks are often downregulated at the cell-cell contact, which is at least in part, a direct result of cadherin activated signaling cascades [10]. In the cell-doublet experiment, E-cadherin and F-actin are homogeneous on the cell-cell contact at the initial stage of the contact formation but diminish in the contact center as the contact matures. This may result from competition between α-catenin and Arp2/3 for the actin binding site [44]. A more recent study suggested that F-actin protrusions are necessary to push cells together to initiate contact [43]. Another experiment showed that, two cells establishing intercellular contact, Rac1 is prominent at the contact edge, which induces the lamellipodia in this area [33]. Meanwhile, Arp2/3-dependent branching network and the formin mDia1-dependent F-actin bundle formation compete for actin pool, which dominate the early and later phases respectively [45–47]. All of these effects of F-actin protrusion, competition between α-catenin and Arp2/3, or competition between formin and Arp2/3 might be interpreted as a singular cause for the ring distribution. But most likely, these mechanisms function simultaneously, including the one suggested by our model: cortical tension induced reduced E-cadherin diffusivity locally and the positive feedback loop between E-cadherin and F-actin.

F-actin affects cadherin diffusion physically by anchoring and building “fences”. Tailless E-cadherin that is not affected by actin microfilaments can form larger but less stable clusters [17, 28, 38]. Although we focus on anchoring, it is worth noting that actin microfilament can fence cadherins and prevent them from reaching each other, which is another important parameter to be considered. A recent study showed that E-cadherin clusters on the lateral junction and the apical junction are different in size and distribution [17]. This result might also imply that the actin microfilament structure can regulate E-cadherin clustering since F-actin is structured in bundles on the apical junction and branched network on the lateral junction.

We have shown using mechanistic modelling how spatially varying mechanical force can trigger a positive feedback loop between the F-actin cortical tension and E-cadherin clustering that subsequently re-organises and re-models the F-actin cortical network and E-cadherin distribution at the cell-cell interface. We predict that different levels of tension across a tissue monolayer will generate heterogeneous E-cadherin distributions at the cell-cell interface. As mechanical tension at E-cadherin junctions are known to regulate intracellular signaling and mechanics, regulation of E-cadherin distribution may affect cell and tissue differentiation. We speculate one could harness the feedback between mechanical force, E-cadherin dynamics and cortical F-actin dynamics to engineer new tissues, or control cancer metastasis.

## Acknowledgments

We thank Jiawen Chen, Xumei Gao, Virgile Viasnoff for helpful discussions.

## Data sharing plan

The model used in this work and the code used for analysis are shared on the GitHub. https://github.com/QILINY/Research_unimelb

## Notes

https://github.com/QILINY/Research_unimelb

### Competing Interest Statement

The authors have declared no competing interest.

