## supplemental materials for "Cortical Tension Initiates the Positive Feedback Loop Between E-cadherin and F-actin"

**This PDF file includes:**

Supplementary text
Figures S1 to S8
Tables S1 to S2
Legends for Movies S1 to S6
SI References

**Other supplementary materials for this manuscript include the following:**

Movies S1 to S6

### Supplementary Information Text

#### Reaction diffusion model of the coupled model

Interactions between the actomyosin network and E-cadherin molecules affect the dynamics of each other. Thus, a mathematical model of the actomyosin network is necessary to study the maturation of the intercellular junction. We created a minimal reaction-diffusion model that only includes key reactions that are necessary to develop the model of intercellular contact formation.

**(1).** E-cadherin homophilic interaction (trans dimerization) upregulates Src, and Src upregulates Rac. [1]

$$Ecad_{trans} + Rac \xrightarrow{k_1} Rac^* + Ecad_{trans}$$

**(2).** Rac stimulates polymerization

$$Rac^* + actin \xrightarrow{k_2} Rac^* + actin^*$$

**(3).** Rac autoactivation and deactivation

$$Rac \underset{k_{3r}}{\overset{k_3}{\longleftrightarrow}} Rac^*$$

**(4).** Actin depolymerization

$$actin^* \xrightarrow{k_4} actin$$

**(5).** Bond formation between E-cadherin and F-actin is mediated through many other proteins such as α-catenin which acts as a catch-slip bond [2]. This bond is simplified as a direct bond between E-cadherin and F-actin. k5r is a function of the force $F_\alpha$ sustained in each single bond between E-cadherin and F-actin, which is discussed in detail below.

$$Ecad + actin^* \xrightarrow{k_5} (Ecad * actin^*) + actin^*$$

$$Ecad + actin^* \underset{k_{5r}}{\longleftarrow} (Ecad * actin^*)$$

$$k_{5r} = f(F_\alpha)$$

**(6).** Rac indirectly inhibits RhoA via p190B-RhoGAP [3].

$$Rac^* + RhoA^* \xrightarrow{k_6} Rac^* + RhoA$$

**(7).** RhoA autoactivation and deactivation

$$RhoA \underset{k_{7r}}{\overset{k_7}{\longleftrightarrow}} RhoA^*$$

**(8).** RhoA activates myosin

$$RhoA^* + myo \xrightarrow{k_8} RhoA^* + myo^*$$

**(9).** Phosphorylated myosin binds F-actin [4]

$$myo^* + actin^* \xrightarrow{k_9} actin^* + (myo^* * actin^*)$$

$$(myo^* * actin^*) \xrightarrow{k_{9r}} myo$$

**(10).** Jasplakinolide (Jasp) induces actin polymerization, an optional parameter in Fig. 6 D in the main text to recapitulate experimental observations.

$$actin \xrightarrow{k_{10}} actin^*$$

### Coupling between the lattice-based model and the continuum model

We use a lattice-based model to simulate E-cadherin dynamics, and a continuum model to simulate the actomyosin dynamics and the interaction between the E-cadherin and the actomyosin network. Due to the discrepancy between the lattice size and the mesh size, a coupling between two models is required.

Firstly, using the package *Triangle*, we generated a triangular mesh, 'Mesh_1', with a radius of 0.5 um to represent the intercellular contact area (Fig. S5 a). For each element in the continuum mesh, we find all the lattice points in that element, according to their positions (Fig. S5 b). Then we map the concentration of E-cadherin in current element to the incenter of each element, and all incenters are used as new nodes to create a mesh, 'Mesh_2' (Fig. S5 c,d). Thus, each element in 'Mesh_1' has a corresponding node in 'Mesh_2'. Concentration of E-cadherin in the element of 'Mesh_1' is calculated based on the information from the lattice-based model and is reported to the corresponding incenter node in the 'Mesh_2', and all reaction-diffusion PDEs will be solved on the 'Mesh_2'. Meanwhile, the mobile state of E-cadherin in the lattice-based model is determined by the concentration of actin in the continuum model.

### Force analysis in the intercellular contact

Contraction of the cytoskeleton beneath the cell membrane is the main factor of the sustained tension at the cell surface, working similar to the surface tension, minimizing the surface area of a cell. When two adhesive cells are brought together, trans-dimerization induced signal cascades diminish the cytoskeleton beneath the contact membranes, and lower the cortical tension at the contact area, until the force equilibrium is attained again. During this process, cortical tension at the free surface, β, is balanced by the remaining cortical tension at the contact area $\beta_{cont}$, adhesion tension Γ, and tension sustained by the E-cadherin trans-dimer λ [5] (Fig. S6).

$$\sum F_x = \beta * \cos(\theta) - \beta_{\mathrm{cont}} - \frac{\Gamma}{2} = 0$$
$$\sum F_y = \beta * \sin(\theta) - \lambda = 0$$

The cortical tension used in the model is 4000 pN/μm, which is then converted to cross-sectional stress of 8000 pN/μm$^2$ by dividing the cortical thickness of a normal size cell, 0.5 μm. We multiply 8000 pN/μm$^2$ by the total cross-sectional area of the cortex on in our model $\pi * (r_{cont}^2 - r_{cent}^2)$, or π*(0.5$^2$ - 0.45$^2$ μm$^2$), to get the total intercellular forces applied on the contact rim, 1194 pN. $r_{cont}$ is radius of the cell-cell contact; $r_{cent}$ is the radius of the contact center, contact area excluding the edge; w is the thickness of the cortex or the width of the contact edge. The initial myo*-actin* concentration is used to ensure the cortical force is balanced by the force generated by myosin on the contact, assuming that all contractility comes from myo*-actin*.

We can calculate $\beta_{cont}$ by multiplying the total number of acto-myosin complexes (myo*-actin*) on the contact with the force per myosin head (4 pN) [6, 7] and accounting for 2 heads being present in each myosin molecule. Total number of myosins is then the summation of the product of the concentration of acto-myosin ([myo*-actin*]) complexes in each mesh element times the element area:

$$\beta_{\mathrm{cont}} = 4\mathrm{pN} * 2 * \sum_{i=1}^{n} ([\mathrm{myo}^{*}-\mathrm{actin}^{*}]_{\mathrm{i}} * \mathrm{Area}_{\mathrm{i}})$$

### E-cadherin/F-actin dissociation rate k5r is governed by α-catenin/F-actin two-state catch-bond

Experiments showed that E-cadherin, β-catenin, α-catenin form a minimal cadherin-catenin complex that binds to the actin cytoskeleton directly in epithelial tissues. Craig D. Buckley and coworkers developed an optical trap-based assay to measure the lifetime of the bond between cadherin-catenin complex and F-actins [2]. They proposed a two-state catch-bond model based on their results, which gives the following relationship between the mean lifetime of the α-catenin/F-actin bond and the force exerted on it (Fig. 7).

Cortical tension also affects the lifetime α-catenin/F-actin bond. It was found that the angle ε between F-actin and α-catenin should be in the range between 11˚ and 127˚ [8] (Fig. S8). We use the mean value of the angle 69˚ in our model. Therefore, we can calculate the force sustained in the α-catenin/F-actin bond, $F_\alpha$, using the following formula:

$$F_\alpha = \frac{\lambda}{N_{\mathrm{Ecad*actin^*}}^{edge}\sin(\varepsilon)}$$

$N_{\mathrm{Ecad*actin^*}}^{edge}$ is the number of E-cadherin molecules that are involved in the F-actin bond on the contact edge. With $F_\alpha$, we can determine the dissociation rate k5r with the following relationship:

$$k_{5r} = \frac{1}{Mean\ Lifetime(F_\alpha)}$$

Details of the mean lifetime function can be found in the Craig D. Buckley's work [2].

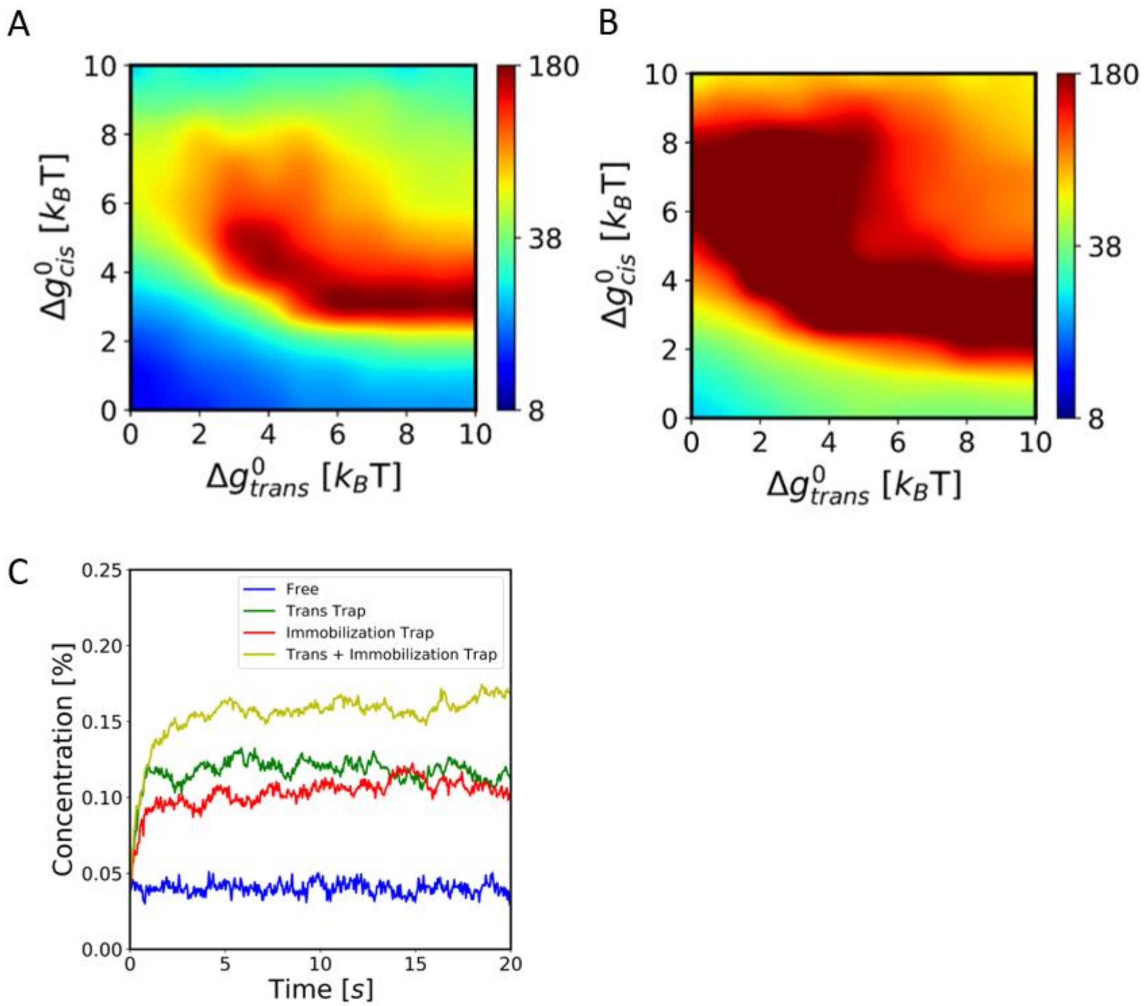


**Fig. S1.** Cluster size analysis with a higher E-cadherin concentration Maximum size of clusters with E-cadherin concentration (A) 0.08, (B) 0.16. Trans-dimers can be formed everywhere. All E-cadherin monomers and trans-dimers can diffuse freely on all the lattices. (C) Concentration of E-cadherin in the central 20 x 20 lattice site for simulations in Figure 2 A-D, $\boldsymbol{\Delta g^0_{trans} = 6.0\ \mathrm{kT}}$, $\boldsymbol{\Delta g^0_{cis} = 2.0\ \mathrm{kT}}$, cadherin concentration = 0.04

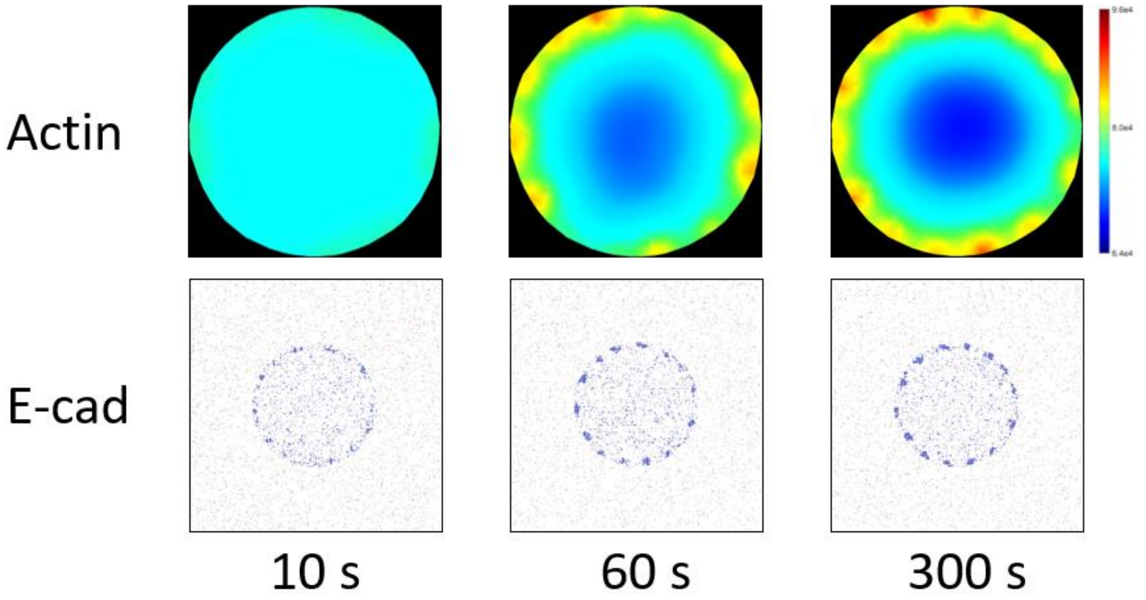


**Fig. S2.** Simulation of the intercellular contact formation of a cell doublet. $\boldsymbol{\Delta g^{0}_{trans} = 6.0\ \mathrm{kT}}$, $\boldsymbol{\Delta g^{0}_{cis} = 2.0\ \mathrm{kT}}$, cadherin concentration = 0.04, $\boldsymbol{k_1 = 5 * 10^{-6}\mu m^2 s^{-1}}$, $\boldsymbol{k_5 = 0.03\mu m^2 s^{-1}}$

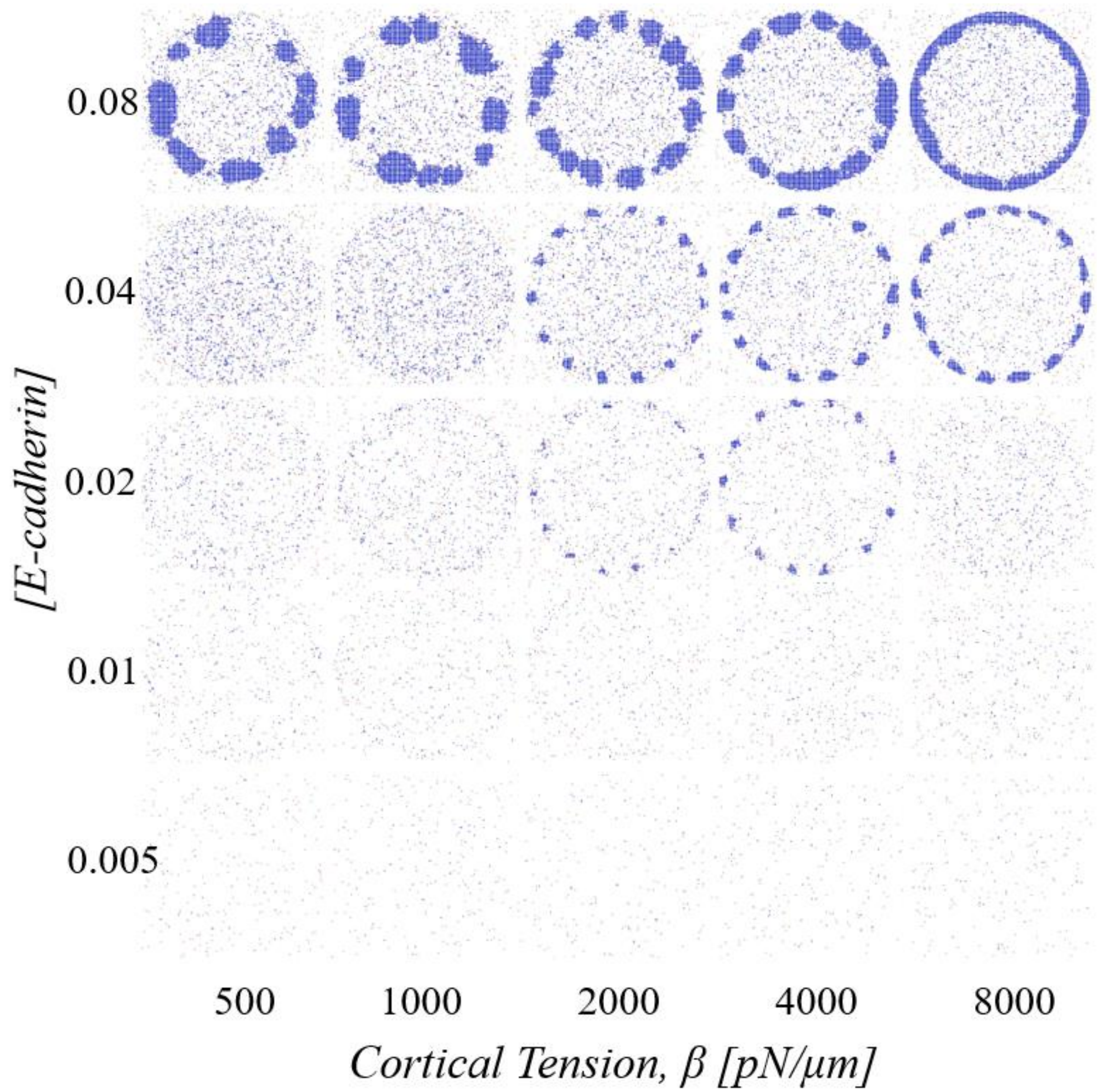


**Fig. S3.** Snapshots of the E-cadherin lattice-based model at 300s with a wider range of E-cadherin concentration and Cortical tension. Only the circular contact region is shown.

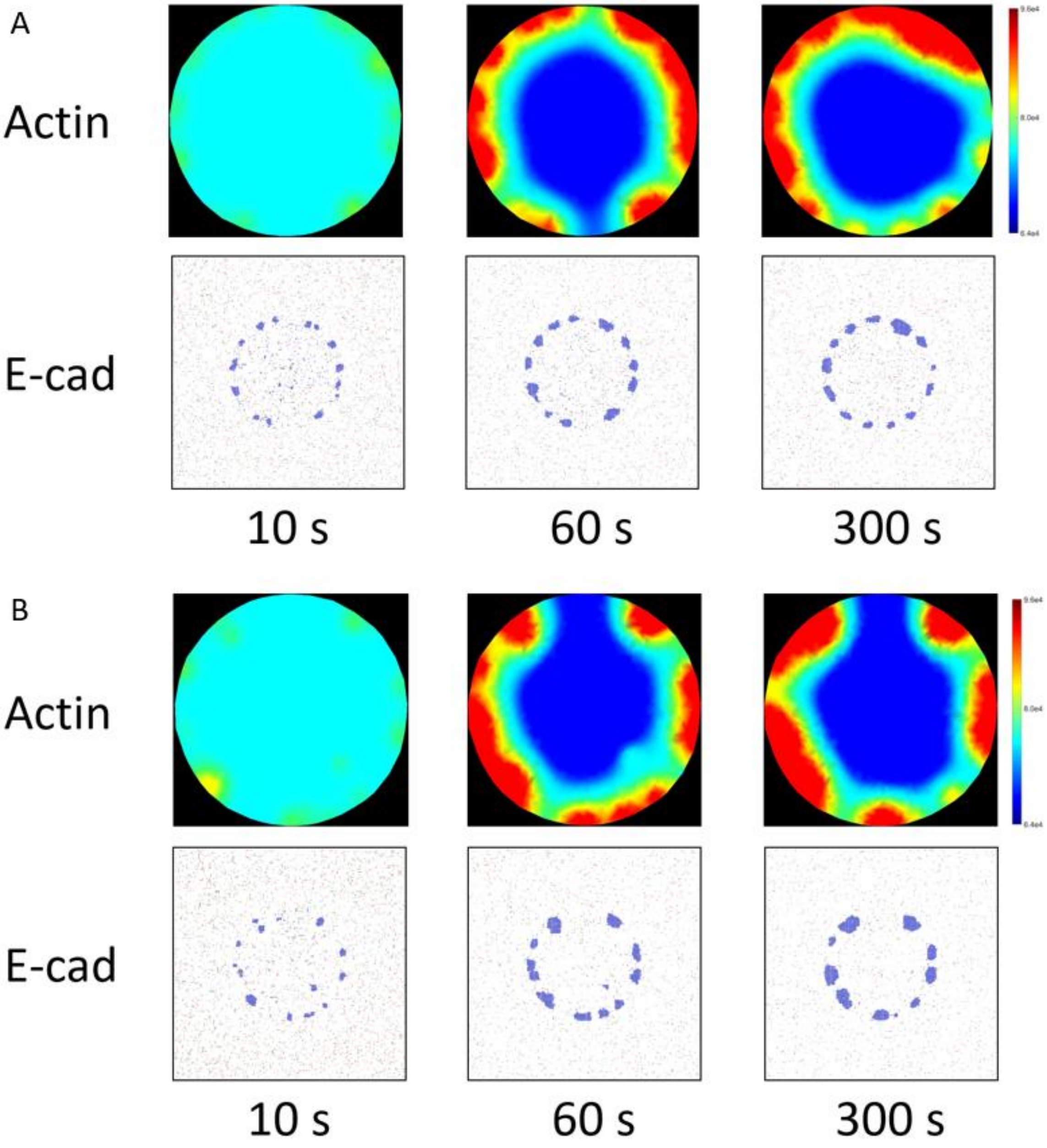
A
Actin
E-cad
10 s
60 s
300 s
B
Actin
E-cad
10 s
60 s
300 s

**Fig. S4.** E-cadherin molecules and F-actin distribution with other combinations of trans and cis binding affinities. E-cadherin molecules concentration = 0.04 As suggested by the experimental results, the binding affinity of trans should be in the range of 3.5-5.5 kT. The binding affinity of cis should be lower. However, the value of cis binding affinity is arguable, because some studies suggest that the cis binding affinity should be higher if E-cads form trans dimers. Here we show the actin and E-cadherin molecules distribution with impact from cortical tension and other value of $\boldsymbol{\Delta g_{trans}^{0}}$and $\boldsymbol{\Delta g_{cis}^{0}}$. It's noticeable that E-cadherin molecules and F-actin can still form a ring-like structure, but the E-cadherin clusters are larger and more stable. This result means that the impact from cortical tension does not depend on the trans and cis binding affinity. (A) $\boldsymbol{\Delta g_{trans}^{0}} = \mathbf{5.0\ kT}$, $\boldsymbol{\Delta g_{cis}^{0}} = \mathbf{3.0\ kT}$ (B) $\boldsymbol{\Delta g_{trans}^{0}} = \mathbf{4.0\ kT}$, $\boldsymbol{\Delta g_{cis}^{0}} = \mathbf{4.0\ kT}$, cadherin concentration = 0.04, $\boldsymbol{k_1 = 5 * 10^{-6}\mu m^2 s^{-1}}$, $\boldsymbol{k_5 = 0.03\mu m^2 s^{-1}}$

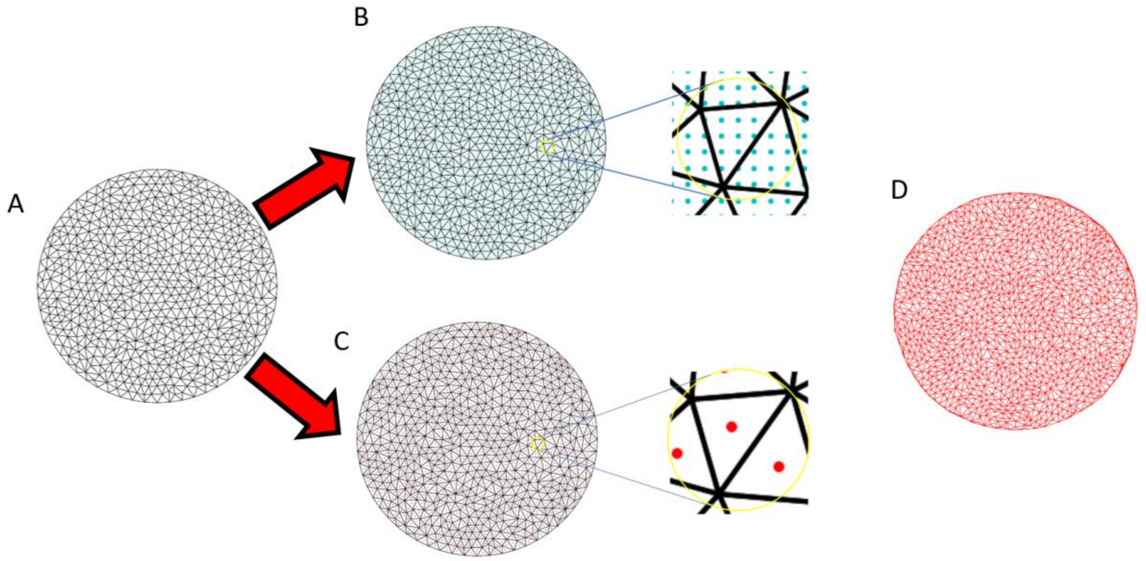


**Fig. S5.** Schematics of the coupling processes between the continuum model and the lattice-based model. (A)The original mesh, ‘Mesh_1’. (B) Correspondence between the original mesh (black) and the lattice points (cyan). (C) Incenters (red) of elements in ‘Mesh_1’. (D) Incenters of elements in ‘Mesh_1’ are used as nodes to generate ‘Mesh_2’. Concentration of E-cadherin molecules is updated to the nodes in ‘Mesh_2’ every time step.

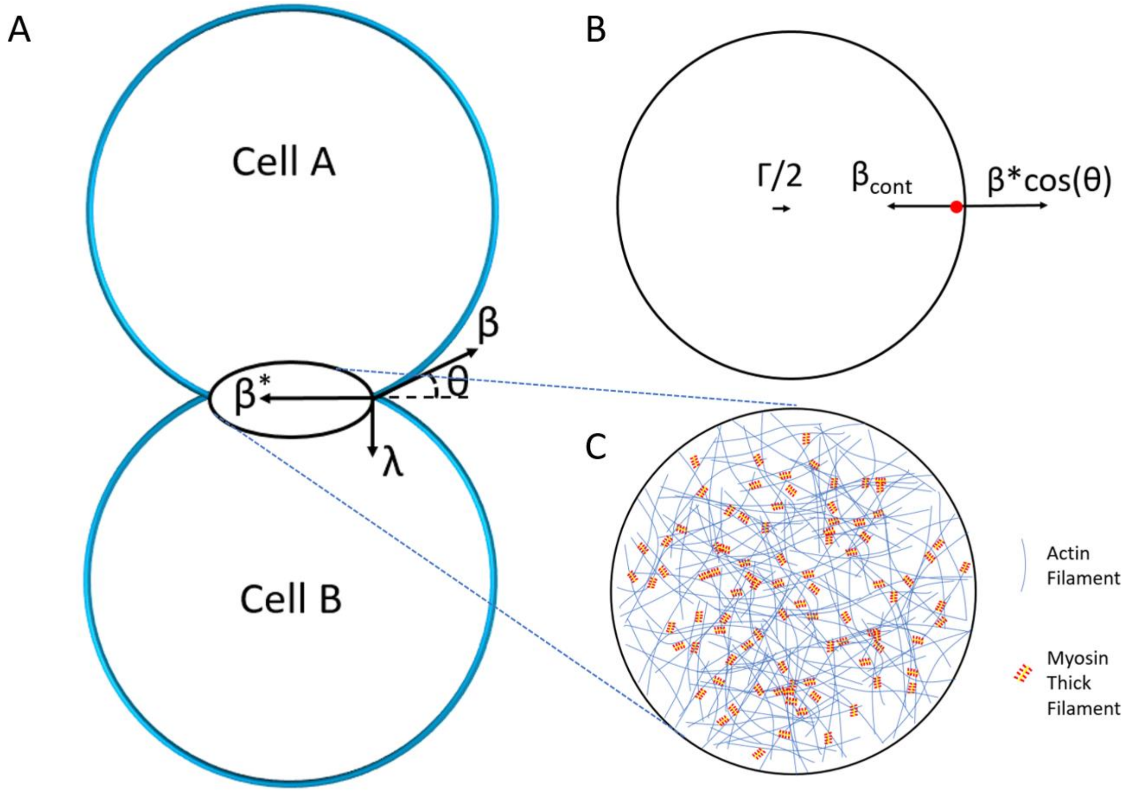


**Fig. S6.** Decomposition of forces on the contact area. (A) Force analysis on the intercellular contact in 3D. (B) Forces parallel to the cell-cell contact. (C) Actomyosin network on the cell-cell contact.

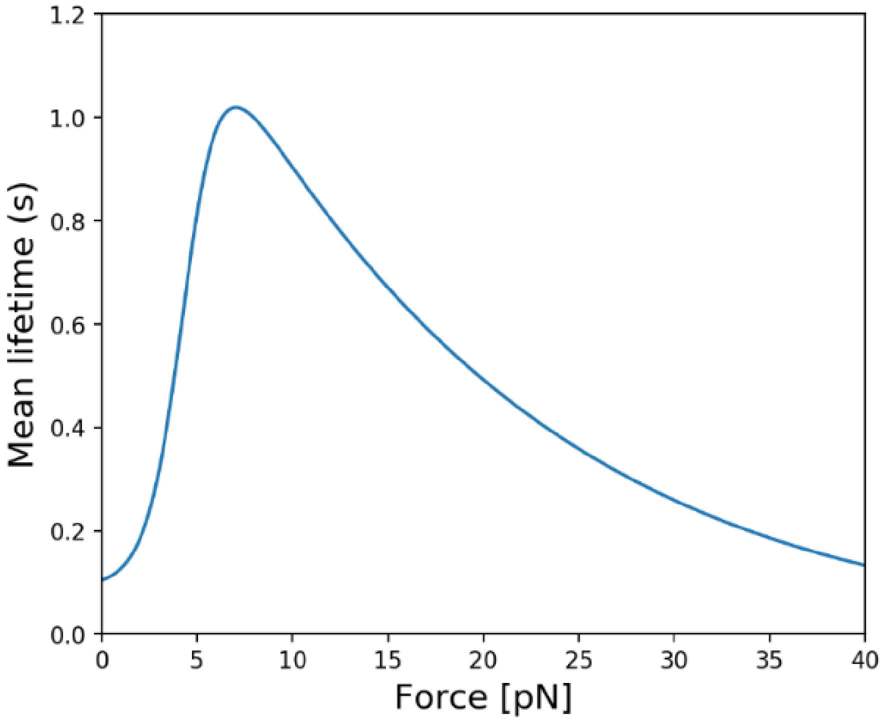


**Fig. S7.** Mean lifetime of the E-cadherin/F-actin bond as a function of force. The binding between E-cadherin and F-actin is simplified using the property of the two states catch-slip bond of α-catenin/F-actin [2].

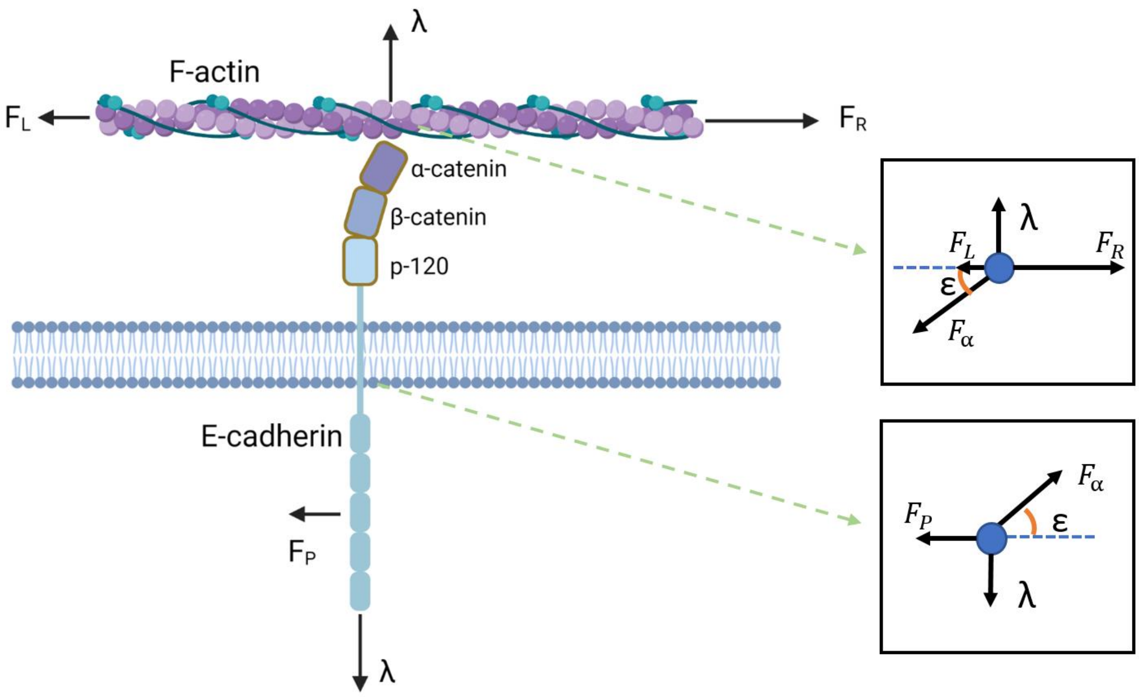


**Fig. S8.** Force analysis of the system that comprises cadherin-catenin complex and F-actin at static state. All external forces are labeled on the figure. Analyses of internal forces are demonstrated in the insets. $\boldsymbol{F_L}$ and $\boldsymbol{F_R}$ are parallel forces applied on the actin cortex; $\boldsymbol{\lambda}$ is the cortical tension transmitted from cell cortex to the E-cadherin trans-dimer; $\boldsymbol{F_P}$ represents any other parallel force applied on the E-cadherin; $\boldsymbol{F_\alpha}$ is the force sustained in the α-catenin/F-actin bond, which affects lifetime of the bond; ε is the angle between α-catenin and F-actin.

| Chemical species | Concentration ($\mu m^{-2}$) | Diffusivity ($\mu m^2 s^{-1}$) | Description | Reference |
|---|---|---|---|---|
| $\boldsymbol{Ecad}$ | * | * | All E-cadherins that are not involved in the bond with F-actin | |
| $\boldsymbol{Ecad_{trans}}$ | * | * | Free E-cadherin trans dimers that are not involved in the bond with F-actin | |
| $\boldsymbol{Ecad * actin^*}$ | * | * | E-cadherin trans dimers that are involved in the bond with F-actin | |
| $\boldsymbol{actin}$ | 5119 | 0.1 | G-actin or actin monomer | [9] |
| $\boldsymbol{actin^*}$ | 2408 | 0 | Actin monomers in the F-actin | [9] |
| $\boldsymbol{Rac}$ | 202.5 | 0.1 | Inactivated Rac | [10] |
| $\boldsymbol{Rac^*}$ | 22.5 | 0.0 | Activated Rac | [10] |
| $\boldsymbol{RhoA}$ | 83.7 | 0.1 | Inactivated RhoA | [10] |
| $\boldsymbol{RhoA^*}$ | 9.3 | 0.0 | Activated RhoA | [10] |
| $\boldsymbol{myo}$ | 2976.67 | 0.1 | Inactivated myosin | |
| $\boldsymbol{myo^*}$ | 0 | 0.1 | Activated myosin | |
| $\boldsymbol{myo^* - actin^*}$ | 190 | 0.0 | Activated myosin that binds to F-actin | |

**Table S1.** Parameters for the Chemical species. Concentrations in mol are converted to 2D assuming a thickness of actin cortex of 0.05 μm, which approximates to the effective thickness of actin that can affect cadherin dynamics [11]. Concentration of acto-myosin complexes ([myo*-actin*]) is computed to ensure that the cortical tension is balanced by the initial contractile force on the contact $\boldsymbol{\beta_{cont}}$. The total myosin concentration is calculated using the duty ratio of 0.06 [9].

* Defined by the Lattice-based model

| Kinetic constants | Value | Description | Reference |
|---|---|---|---|
| k1 | $10^{-7}$ - $10^{-3}$ $\mu m^2 s^{-1}$ | Rac activation rate by Trans dimerization | |
| k2 | 0.0006272 $\mu m^2 s^{-1}$ | Actin polymerization rate activated by Rac | |
| k3 | 0.01 $s^{-1}$ | Rac activation rate | [10] |
| k3r | 0.09 $s^{-1}$ | Rac deactivation rate | [10] |
| k4 | 0.03 $s^{-1}$ | Actin depolymerization rate | [12] |
| k5 | $10^{-5}$ - $10^{-1}$ $\mu m^2 s^{-1}$ | E-cadherin/F-actin binding rate | |
| k5r | ~1-10 $s^{-1}$ | E-cadherin/F-actin dissociation rate | [2] |
| k6 | 0.05 $\mu m^2 s^{-1}$ | Rho inhibition rate by Rac | |
| k7 | 0.136 $s^{-1}$ | Rho activation rate | [10] |
| k7r | 0.1 $s^{-1}$ | Rho deactivation rate | |
| k8 | 0.26 $\mu m^2 s^{-1}$ | Myosin activation rate by Rho | |
| k9 | 0.2 $\mu m^2 s^{-1}$ | Myosin binding rate to actin filaments | |
| k9r | 37.6 $s^{-1}$ | Myosin unbinding rate from F-actin | [13] |
| K10 | 0 or 1 $\mu m^2 s^{-1}$ | Jasp induced actin polymerization | |

**Table S2.** Values of kinetic constants used in the reaction diffusion model. k0 is an optional parameter used in the study of Fig. 6 D of the main text. k1, k5 are key parameters, which are studied in Fig. 3 in the main text. Kinetic constants k3,k3r,k7,k7r are set assuming that the active fraction of Rac and RhoA, Rac*, RhoA*, take 10% of total concentration [10]. Kinetic constants k2, k6, k7r, k8, k9 are adjusted to make sure all species are in an equilibrium state when there is no E-cadherin trans-dimers formation in the model; also to make sure that Rac and Rho are in the reasonable concentration with stimulation of E-cadherin trans-dimer formation [14].

**Movie S1 (separate file).** F-actin distribution overtime of the continuum model, $\boldsymbol{\Delta g_{trans}^{0}} = \mathbf{6.0\ kT}$, $\boldsymbol{\Delta g_{cis}^{0}} = \mathbf{2.0\ Kt}$, $\textbf{cadherin concentration} = \mathbf{0.04}$, $\boldsymbol{k_1} = \mathbf{5 * 10^{-6}}\mu\boldsymbol{m^2 s^{-1}}$, $\boldsymbol{k_5} = \mathbf{0.03}\mu\boldsymbol{m^2 s^{-1}}$

**Movie S2 (separate file).** E-Cadherin distribution overtime of the lattice-based model, $\boldsymbol{\Delta g_{trans}^{0}} = \mathbf{6.0\ kT}$, $\boldsymbol{\Delta g_{cis}^{0}} = \mathbf{2.0\ Kt}$, $\textbf{cadherin concentration} = \mathbf{0.04}$, $\boldsymbol{k_1} = \mathbf{5 * 10^{-6}}\mu\boldsymbol{m^2 s^{-1}}$, $\boldsymbol{k_5} = \mathbf{0.03}\mu\boldsymbol{m^2 s^{-1}}$

**Movie S3 (separate file).** F-actin distribution overtime of the continuum model, $\boldsymbol{\Delta g_{trans}^{0}} = \mathbf{6.0\ kT}$, $\boldsymbol{\Delta g_{cis}^{0}} = \mathbf{2.0\ Kt}$, $\textbf{cadherin concentration} = \mathbf{0.04}$, $\boldsymbol{k_1} = \mathbf{5 * 10^{-6}}\mu\boldsymbol{m^2 s^{-1}}$, $\boldsymbol{k_5} = \mathbf{0.03}\mu\boldsymbol{m^2 s^{-1}}$

**Movie S4 (separate file).** E-Cadherin distribution overtime of the lattice-based model, $\boldsymbol{\Delta g_{trans}^{0}} = \mathbf{6.0\ kT}$, $\boldsymbol{\Delta g_{cis}^{0}} = \mathbf{2.0\ Kt}$, $\textbf{cadherin concentration} = \mathbf{0.04}$, $\boldsymbol{k_1} = \mathbf{5 * 10^{-6}}\mu\boldsymbol{m^2 s^{-1}}$, $\boldsymbol{k_5} = \mathbf{0.03}\mu\boldsymbol{m^2 s^{-1}}$

**Movie S5 (separate file).** F-actin distribution overtime of the continuum model, $\boldsymbol{\Delta g_{trans}^{0}} = \mathbf{6.0\ kT}$, $\boldsymbol{\Delta g_{cis}^{0}} = \mathbf{2.0\ Kt}$, $\textbf{cadherin concentration} = \mathbf{0.04}$, $\boldsymbol{k_1} = \mathbf{5 * 10^{-6}}\mu\boldsymbol{m^2 s^{-1}}$, $\boldsymbol{k_5} = \mathbf{0.03}\mu\boldsymbol{m^2 s^{-1}}$

**Movie S6 (separate file).** E-Cadherin distribution overtime of the lattice-based model, $\boldsymbol{\Delta g_{trans}^{0}} = \mathbf{6.0\ kT}$, $\boldsymbol{\Delta g_{cis}^{0}} = \mathbf{2.0\ Kt}$, $\textbf{cadherin concentration} = \mathbf{0.04}$, $\boldsymbol{k_1} = \mathbf{5 * 10^{-6}}\mu\boldsymbol{m^2 s^{-1}}$, $\boldsymbol{k_5} = \mathbf{0.03}\mu\boldsymbol{m^2 s^{-1}}$
